# Deciphering transcriptional programming during lupin (*Lupinus angustifolius*) seed development using RNA-seq

**DOI:** 10.1101/2024.09.05.611346

**Authors:** Virginia Wainaina, Tina Rathjen, Trijntje Hughes, Annelie Marquardt, Natalie Fletcher, Hayley Casarotto, Meredith McNeil, Kerensa McElroy, Ling-Ling Gao

**Affiliations:** Commonwealth Scientific and Industrial Research Organization, Floreat, WA; Commonwealth Scientific and Industrial Research Organization, Black Mountain, ACT, 2601; Commonwealth Scientific and Industrial Research Organization, St Lucia, QLD, 4067

**Keywords:** *Lupinus angustifolius*, seed development, transcriptome, annotation, differential expression

## Abstract

Lupin (*Lupinus spp.)* seeds are valued for their high protein content (35-40%) for both human and animal consumption. Seed development in crop plants is a critical factor influencing both seed fate and yield, hence, understanding the molecular mechanisms of seed development is essential. This study conducted a transcriptome analysis of Narrow Leaf Lupin (NLL) during seed development stages (3, 6, 9, 12, 15, 18, and 21 days after flowering) to investigate transcriptional dynamics and identify key candidate genes that control seed development. Approximately 357 million sequencing reads were generated from nine samples from leave, flower and seed tissues, enabling the identification of 34,769 expressed genes. The analysis revealed dynamic gene expression, with early stages marked by high metabolic activity and later stages focusing on storage protein synthesis and nutrient reservoir formation. The differential expression patterns of seed storage protein genes, including cupin groups (α, β, γ, and δ conglutins), were notable. The expression of α and β conglutins increased at later stages (15-21 days after flowering), supporting their role in grain filling and nutrient storage. Genes related to quinolizidine alkaloid biosynthesis, such as lysine/ornithine decarboxylase and purine permease transporter 1, showed late expression patterns suggesting alkaloid synthesis and transport during later stages. Many of the well-established transcription factors (TFs) known for their roles in seed development (bHLH, AP2, MYB, ERF, C2H2, NAC, WRKY, and C3H zinc finger families) showed differential expression, thus reinforcing the validity of our findings. These findings lay the groundwork for understanding the genetic and molecular mechanisms of seed development in lupin, contributing to enhanced crop management and breeding programs.

## Introduction

The surge in the global human population, projected to reach 9.7 billion by 2050, stands as one of the foremost challenges confronting our world [1]. Ensuring food and nutritional security for the continuously expanding human population necessitates the integration of underutilized and neglected crops into mainstream agricultural practices. Protein has been identified as the most limiting macronutrient; therefore, sufficient protein quantity and quality will be required to meeting the increasing global food demand which is estimated to double by 2050 [2].

Lupins (*Lupinus spp.* 2n=40) belong to the Fabaceae family, one of the largest and most diverse families of flowering plants. Lupin grain/seed has gained increasing interest due to its excellent source of plant protein (30-40%), dietary carbohydrate (40%), oil (6%), fibre, numerous minerals, vitamins, and other health-friendly ingredients [3, 4]. [5] Nutrient accumulation processes are regulated by a multitude of molecular activities that occur in a simultaneous coordination [6, 7, 5]. Research focusing on lupin enhancement aims to elucidate the genetic regulations that influence key aspects such as seed composition, grain weight, and seed traits. For instance, a previous transcriptome analysis study identified 16 conglutin genes in the narrow-leafed lupin (NLL) variety Tanjil during seed development, providing insight into the structure and expression of these genes across lupin species at different stages of seed development [5]. However, the broader regulatory framework governing lupin seed development at a global level remains poorly understood. This incomplete knowledge creates significant gaps that impede our ability to fully optimize and enhance the molecular basis of lupin crops.

Exploration of key candidate genes for seed development that are involved in seed size/weight and nutrient accumulation has been reported as significant in legumes [8, 9]. Research into the development of seeds in legumes such as soybean [10, 11, 12], chickpea [6] and model plants *Arabidopsis thaliana* [13], *Medicago truncatula* [14], and *Lotus japonica* [15] has shown that the process is governed by regulatory genes involved in the biosynthesis of important nutrients like protein, carbohydrates and antioxidants. A study on *Arabidopsis* revealed that most genes involved in seed development are expressed across all developmental stages. However, each stage also possesses a distinct subset of genes uniquely active during specific developmental phases [13]. Also, a study investigating gene expression in chickpeas during seed development identified significant enrichment of genes in functional categories such as catalytic activity, binding, cellular processes, metabolic processes, and cellular components including cells and membranes [16]. These findings strongly suggest that seed development is a genetically programmed process, potentially linked to elevated metabolic and catalytic activities within cellular and membrane structures. However, despite various transcriptome analyses in lupin [17, 5], and a draft genome assembled [18], none of this research has focused specifically on global regulatory networks of gene expression during seed development.

Global transcriptomes have been shown to play a crucial role in identifying key genes and regulatory networks during seed development in various plant species. For example, transcriptome studies during seed development in legumes such as lentil [19], soybean [10], peanut [20, 7] and chickpea [8], and cereals including rice [21], wheat [22] and maize [23] have demonstrated the power of global transcriptome analysis in unravelling the genetic basis of seed development. These studies demonstrate that by profiling gene expression patterns, researchers can pinpoint genes involved in crucial processes like embryo formation, endosperm development, and seed coat composition. Additionally, transcriptome analysis enables the identification of regulatory networks and transcription factors that govern seed development as shown in the model plants *Arabidopsi*s *thaliana* [13], *Brassica juncea* [24], and *Medicago sativa* [25]. These comprehensive approaches not only enhance our understanding of seed development mechanisms, but researchers identify key regulatory hubs and potential targets for genetic improvement of seed traits. Application of this global profiling leads to more effective strategies for enhancing seed development and crop performance of important crop species such as lupin.

Sequence-based transcriptome profiling (RNA-seq) has emerged as a powerful approach for the identification of differentially expressed genes across conditions, aiding in the exploration of molecular functions [26, 27]. While previous research studies have extensively employed high throughput techniques to investigate seed development in various plant species such as *Arabidopsis* [13], wheat [28], rice [29], peanut [9], soybean [30], chickpea [6] and *Medicago* [31], exploration in the space of lupin is lacking. In this study, we apply RNA-seq to investigate gene expression of progressively developing seed at 3, 9, 12, 15, 18, and 21 days after flowering (DAF), herein depicted at stages A, B, C, D, E, F and G, respectively, to extend our knowledge of the molecular mechanisms that control seed development and nutrient accumulation. We uncovered a variety of gene activities during the dynamic processes of seed development and identified metabolic pathways, and transcription factors associated with seed development. The results provide a global view of gene activities essential to the development processes in lupin, providing insight into the regulatory networks that govern the lupin seed developmental process. The comprehensive dataset of lupin seed transcriptomes serves as a valuable resource for future studies, enabling researchers to explore new aspects of lupin biology and potentially discover novel genes and pathways.

## Material and Methods

### Plant Material

The lupin cultivar, Tanjil, was cultivated under glasshouse conditions. Seeds were germinated and grown in temperature-controlled growth cabinets set to 22°C during a 16-hour day and 20°C during an 8-hour night. Buds were tagged and dated for subsequent harvesting of developing seeds. Flowers and pods, ranging from young to mature, were sampled at intervals of 1-3 DAF (days after flowering), 4-6 DAF, 7-9 DAF, 10-12 DAF, 13-15 DAF, 16-18 DAF, and 19-21 DAF, corresponding to stages A, B, C, D, E, F, and G, respectively. For this study, DAF was determined as the day the flower fully opened with petals completely extended. Seeds from each tagged set were manually separated from their pods, immediately frozen in liquid nitrogen, and stored at -80°C. For RNA extraction, more than one developing seed per plant were combined to form a single biological replicate, with three such replicates used for each developmental stage. Throughout the manuscript, samples are represented as seed stages A, B, C, D, E, F, and G.

### RNA isolation and transcriptome sequencing

Approximately 150 milligrams of plant material from each tissue type were ground into a fine powder using liquid nitrogen. Total RNA was isolated from foliar and seed tissues using TRIzol reagent (Invitrogen, CA, USA) following the manufacturer’s protocol as described by [17]. Before library construction, total RNA was treated with RNase-free DNase I (New England Biolabs, Ipswich, MA, USA) to remove any genomic DNA contamination. RNA samples with integrity values >1.8 were included in the study and quantified using a Nanodrop. Additionally, RNA integrity was verified by resolving a portion of each sample on a denatured agarose gel and assessing the intensity of the 28S and 18S rRNA bands.

### RNA sequencing and data pre-processing

Two micrograms of RNA were sent to the Australian Genome Research Facility (AGRF) in Melbourne, Australia, for library preparation and sequencing. Libraries were prepared using the TruSeq mRNA Library Prep Kit with polyA selection and unique dual indexing. Sequencing was conducted on the Illumina NovaSeq 6000 platform, producing 150-nucleotide-long paired-end reads. The raw reads were subjected to quality filtering using FASTP version 0.23.4-3 to evaluate quality control (QC) metrics, including base quality distribution, and to trim adapters. Following data processing, the raw reads were converted into clean reads.

### Differential gene expression and tissue specificity analysis

These clean reads were aligned and mapped to the reference lupin genome, *L. angustifolius* v.2 genome, [18] using STAR aligner v1.9.0 [33] with default settings. The alignment file from STAR for each sample along with reference genome GTF was used to perform reference annotation-based transcript to assemble genes.

The raw counts per million obtained thereof were normalized using the variance stabilizing transformation (vst) as implemented in DESeq2 v3.16, Heidelberg, Germany [34] in R language [35], with a p-value cut-off of <0.05 after adjustment for multiple testing with the Benjamin Hochberg method. The seed stages were each used as experimental data; the Log_2_ fold change (Log_2_FC) differences in expression data reflect a change occurring in the seed stages relative to stage A (3 DAF). Transcripts with a p-value < 0.05 and |log2FC| ≥ 2 (both up-regulated and down-regulated) were designated as significant differentially expressed genes (DEGs).

K-means clustering analysis of DEGs using k-means function in R, where k=5 within the cluster package by Euclidean distance. Heat maps were drawn using the Pheatmap package in R.

### Gene annotations, GO term, and pathway assignment

For Gene Ontology (GO) enrichment and pathway analysis, each lupin transcript was aligned to the non-redundant (nr) database at NCBI using BLAST with (e-value ≤ 1e-5). The best Arabidopsis thaliana hit corresponding to each lupin transcript was then identified. This approach allowed for the annotation and subsequent functional analysis of lupin transcripts based on their similarity to well-characterized Arabidopsis genes. Gene ontology analysis (GO) is a common method for annotating gene products; it can also decide the defining biological characteristics of high-throughput genomic data. The Kyoto Encyclopedia of Genes and Genomes (KEGG) is a genetic function database that links functional information [37] [38].

### Identification of Transcription Factors

Lupin TFs were identified by aligning sequences against the Plant TF database (Plant TFDB v5.0, https://planttfdb.gao-lab.org/) (E-value 1-5) [39] to annotate and assign them to a TF family.

### Determination of transcript abundance

To elucidate the dynamics of seed development, the uniquely mapped reads were subsequently employed to assess gene expression abundance, normalized as counts per million (CPM). Unigenes with a count value ≥ 1 in at least one seed tissue were considered transcriptionally expressed.

### Real-time quantitative polymerase chain reaction

To validate the accuracy of the RNA-seq data, we performed reverse transcription quantitative polymerase chain reactions (RT-qPCRs) (qRT-PCR) on 10 randomly selected DEGs carried out using the Biorad CFX384 Real-Time PCR system. RNA samples used for qPCR validation were the same samples used for Illumina sequencing. According to the manufacturer’s instruction, the RNA samples were reverse transcribed using the First-Strand cDNA Synthesis Kit for RT-PCR (AMV) (NEB Inc. Ipswitch, MA). The cDNA samples were then diluted 1:20 before qPCR. The gene-specific primers were designed using Genious Prime software version 2024.0.5 (http://www.geneious.com/prime/) with default parameters for qPCR. qPCR reactions were performed using iTaq Univer SYBR Green Master Mix for qPCR with the following cycling conditions: 95 °C for 2 min, then 95 °C for 15 s, 60 °C for 30 s and 72 °C for 30 s, repeated 40 cycles, The lupin TUBULIN gene (GenBank accession: XM_019602599) served as an internal reference, while a negative control reaction utilized water in place of cDNA. Three biological replicates were used, and two technical replicates per biological replicate to determine the expression levels for the ten genes relative β-tubulin. Relative quantification (RQ) values were calculated following the 2^-Δ Δ Ct^ method [40].

## Results

### Transcriptome data profiles

A comprehensive understanding of the biological mechanisms of seed maturation and the accrual of reserves in lupin seeds is pivotal for enhancing seed quality and maximizing yield potential. A time course transcriptomic profiling of seed development at seven sequential times to 3, 6, 9, 12, 15, 18, and 21 days after flowering (DAF), corresponding to A, B, C, D, E, F and G, respectively, was performed to obtain a global view of the regulatory mechanism underlying this complex development process. In total, approximately 357 million sequencing reads were produced from the entirety of sampled seeds, yielding an average of 39 million reads per individual sample, as detailed in Table 1. On average, following the trimming of low-quality reads, 97.43% of reads were retained and utilized for read alignment.

**Table 1:** Summary of the Illumina sequence reads generated by RNA-seq obtained from the seven developmental stages. The value for each stage is the average of the three biological replicates.

| Tissue/Time point | Raw reads | Reads after trimming | Uniquely mapped reads |
| --- | --- | --- | --- |
| Leaf | 44053995 | 43720574 | 39523597 (90.3%) |
| 3 DAF | 40522151 | 40311607 | 36036761(89.4%) |
| 6 DAF | 37985579 | 37813103 | 34033213(90.1%) |
| 9 DAF | 35524040 | 35349547 | 31824779(90%) |
| 12 DAF | 47513874 | 47240923 | 43189579(91.3%) |
| 15 DAF | 35580435 | 35397878 | 32346639(91.3%) |
| 18 DAF | 38503036 | 38313008 | 34778492(90.7%) |
| 21 DAF | 35812496 | 35648166 | 307748779(85.8%) |

The PCA and clustering dendrogram analysis indicated that gene expression levels among the replicated samples within each timepoint were highly correlated (Fig 1), indicating gene expression specificity in each of the time points. Two primary clusters showed the grouping of the samples by the developmental stage. The first cluster represents early stages A, B, C, and D, (representing 3, 6, 9, and 12 DAF) as the early developmental stage and the second cluster comprised the staged E, F, and G (representing 15, 18, and 21 DAF) the late developmental stage.

**Fig 1.**
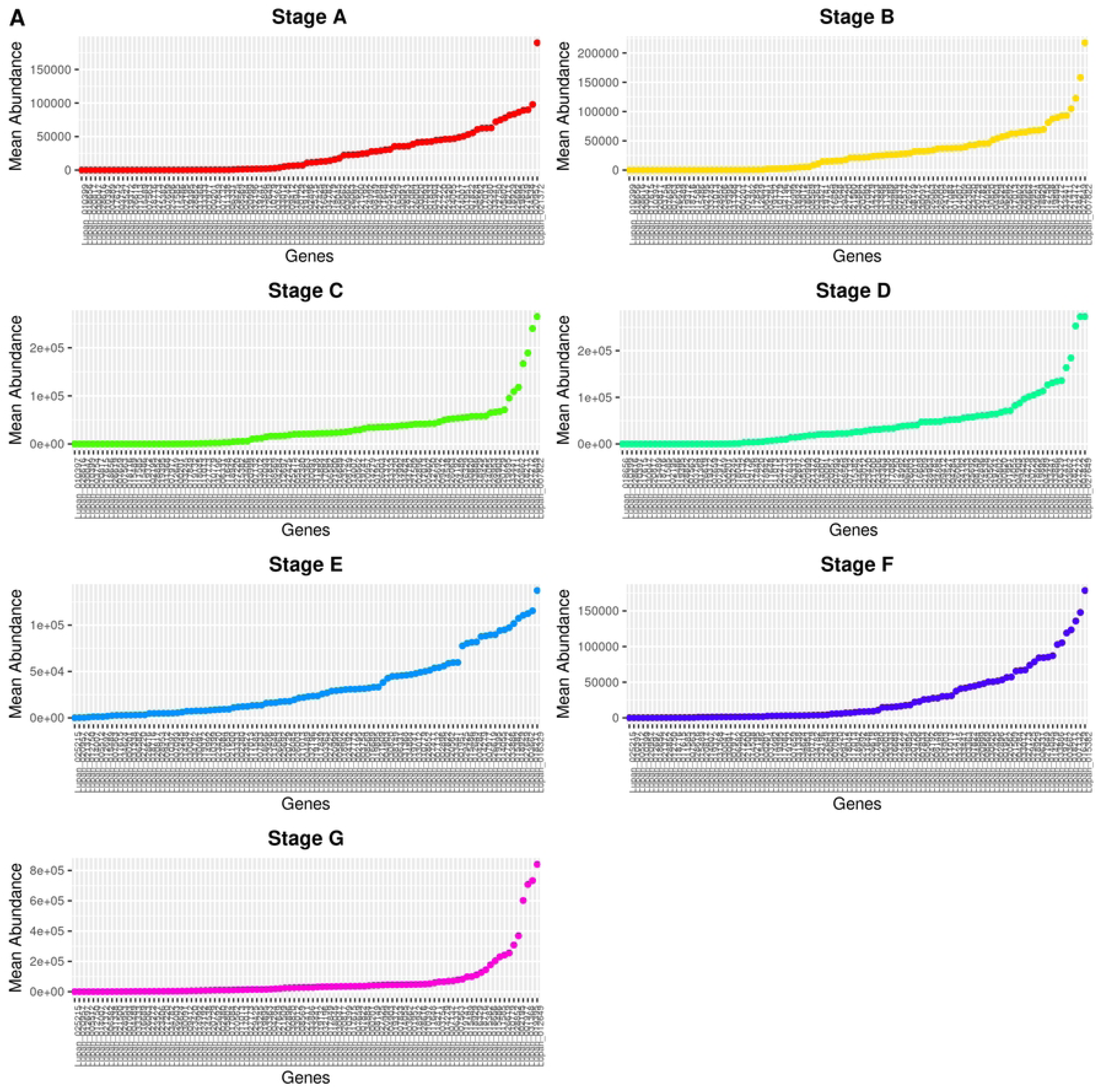
Principal Component Analysis (PCA) of the collected samples across seven stages of seed development, as well as flower and leaf tissues.

### Transcript Abundance and Expression Dynamics in Developing Lupin Seed

Our analysis revealed that, on average, 34,649 gene models, constituting 90.07% of the total 38,466 gene models found in the lupin reference genome, exhibited expression in developing seeds. We conducted an abundance analysis across the developmental stages to identify genes crucial for maintaining overall cellular function, characterized by high abundance. We focused on the top 100 genes based on mean abundance to pinpoint those with the highest average expression levels across stages (Fig 2A). We further subset genes consistently expressed at high levels and analysed the top genes by variance to identify those showing the greatest variability in expression across stages. (Fig S1 A and B).

**Fig 2.**
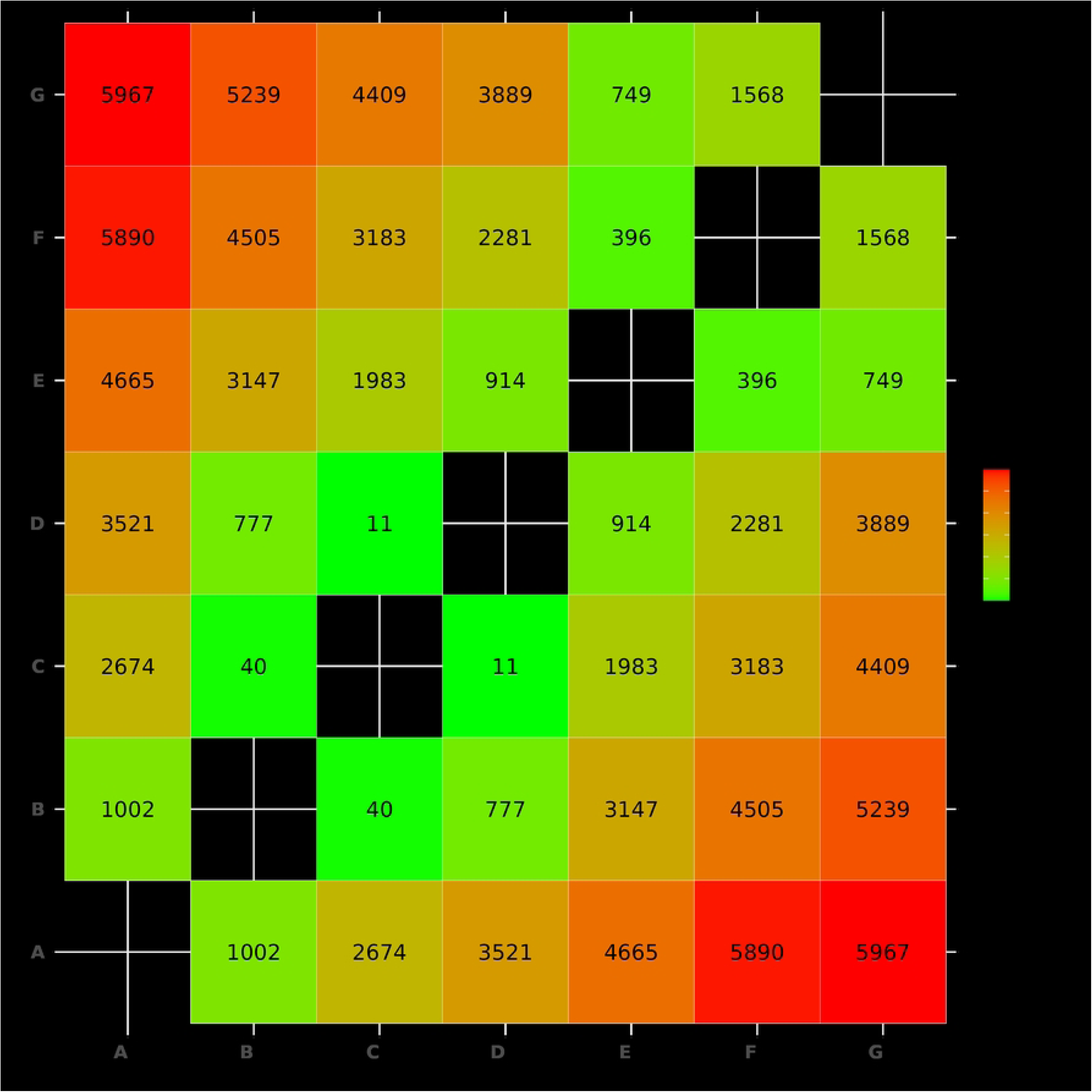
The mean abundance across all stages for the top 100 genes with the highest mean abundance during seed development.

### Highly abundant expressed genes

Among the most highly abundant genes, we identified the top 20 genes exhibiting high abundance across various stages (Fig S1A). For example, Lupan_003019 (Glyoxalase) abundance levels fluctuated across the stages, with a general decrease from stage A to F and a subsequent increase in stage G, reaching its peak in stage G (30,317.67). Lupan_003505 (Adenosylhomocysteinase-like) showed its lowest abundance in stage G (21,926.33) and the highest in stage D (48,713). Lupan_005830 (Tubulin) exhibited significant variations, with the highest abundance in stage E (67,584) and the lowest in stage C (35,031.67). Meanwhile, Lupan_006539 (RNA recognition) displayed its highest abundance in stage A (28,390.67) and the lowest in stage D (20,387.67). The Lupan_008439 (Cobalamin-independent synthase) gene peaked in stage F (40,983.33) and had the lowest abundance in stage B (21,211). Notably, Lupan_011347 (Ubiquitin family) showed the highest abundance in stage D (44,821) and the lowest in stage G (19,676.67). Similarly, Lupan_011701 (Elongation factor) had the highest abundance in stage F (36,935.67) and the lowest in stage G (16,798.67). The Lupan_012244 (Calreticulin) gene displayed the highest abundance in stage D (53,313) and the lowest in stage G (27,709.67). Lupan_014985 (RNA recognition motif domain) exhibited its highest abundance in stage A (48,861) and the lowest in stage D (23,446.33). Lupan_027887 (Heat shock protein) showed its highest abundance in stage F (41,461) and the lowest in stage B (25,351.67).

### Variance in expression abundance

We identified top 20 genes exhibiting high variance in abundance across various stages, indicating their dynamic regulation throughout the different stages (Fig 2C). For example, Lupan_000201 (cupin) showed a significant increase in abundance from stage E (2,869.67) to stage G (39,293.67). Lupan_003754 (Cytochrome P450) had very low abundance in the early stages (A to D), but a dramatic increase in stages E (31,283) and G (65,294). Similarly, Lupan_005512 (Gibberellin regulated protein) exhibited a peak in abundance at stage A (30,142.67) and a rapid decline by stage F (51.33).

Lupan_007093 (Annexin repeat) showed high abundance at stage A (41,732.67), which decreased significantly to stage G (1,421.67). Lupan_007563 (Lipoxygenase) had low abundance in the early stages but peaked dramatically in stage G (77,383). Lupan_007659 (Hypothetical protein) also showed a sharp increase in abundance from stage E (9,244.33) to stage G (35,689.67). Lupan_010397 (cupin) displayed very low abundance in early stages, with a significant increase in stage E (4,926.33) and a peak in stage G (49,961.67).

Lupan_010399 (cupin family) showed minimal abundance from stages A to D, with a sharp increase in stage E (8,217.33) and a peak in stage G (35,127.33). Lupan_015489 (Enoyl-(Acyl carrier protein) reductase) had low abundance in early stages but showed a dramatic increase in stage G (98,620.33). Lupan_017615 (Gamma-thionin family) also had low early-stage abundance and a sharp increase in stage G (35,630.33).

Lupan_017849 (Sucrose synthase) showed increasing abundance from stages A (329.67) to E (23,582.33), peaking in stage G (33,054.33). Lupan_018942 (Ribulose bisphosphate carboxylase) showed gradual increases in abundance, peaking in stage G (47,921). Lupan_019372 (cupin) exhibited low abundance in early stages, with a significant peak in stage G (45,285). Lupan_023527 (Expansin) showed a sharp increase from stage B (10,724.33) to stage D (50,387.33) and then declined in stage G (2,528).

Lupan_027220 (Pollen proteins) displayed high abundance in stage A (44,949), which decreased significantly by stage G (2,706.33). Lupan_027773 (Cellulose synthase) showed a peak in stage F (67,103.33), with lower abundance in other stages. Lupan_028616 (Hypothetical protein) had minimal abundance in early stages, peaking in stage G (48,760.67). Lupan_030091 (Xylanase inhibitor) showed high abundance in early stages, peaking in stage A (35,420.33) and declining by stage G (4,015.67). Lupan_031338 (Hypothetical protein) exhibited a peak in stage D (33,009.67), while Lupan_033245 (Auxin associated protein) showed a peak in stage F (37,481).

### Differential gene expression

While highly abundant genes underscore those necessary for maintaining cellular functions, DEGs highlight the genes essential for transitioning between developmental stages. This integrated approach uncovers the complexity of gene regulation during seed development. To comprehensively identify genes involved in developmental processes during seed development, we performed a pairwise comparison of transcriptomic profiles expression profiles among the different developmental stages of seeds. We observed a gradient increase in the number of DEGs between the two neighbouring points as the seed developed, with 1002, 2674, 3521, 4665, 5890 and 5967 in Stages A vs. B, A vs. C, A vs. D, A vs. E, A vs. F, and A vs. G, respectively (Fig 3, and Table 2S). When comparisons were made across non-adjacent time points (over 3 days interval) the number of genes was low at the early stages, with 40 and 11 in B vs. C and C vs. D, respectively and increased in the later stages with D vs. E and E vs. F and E vs. F with 914, 396 and 1568 genes respectively (Fig 3). In total, the pairwise comparison identified 3224 nonredundant DEGs that are required to program lupin seed development from 3 to 21 days of seed development.

**Fig. 3.**
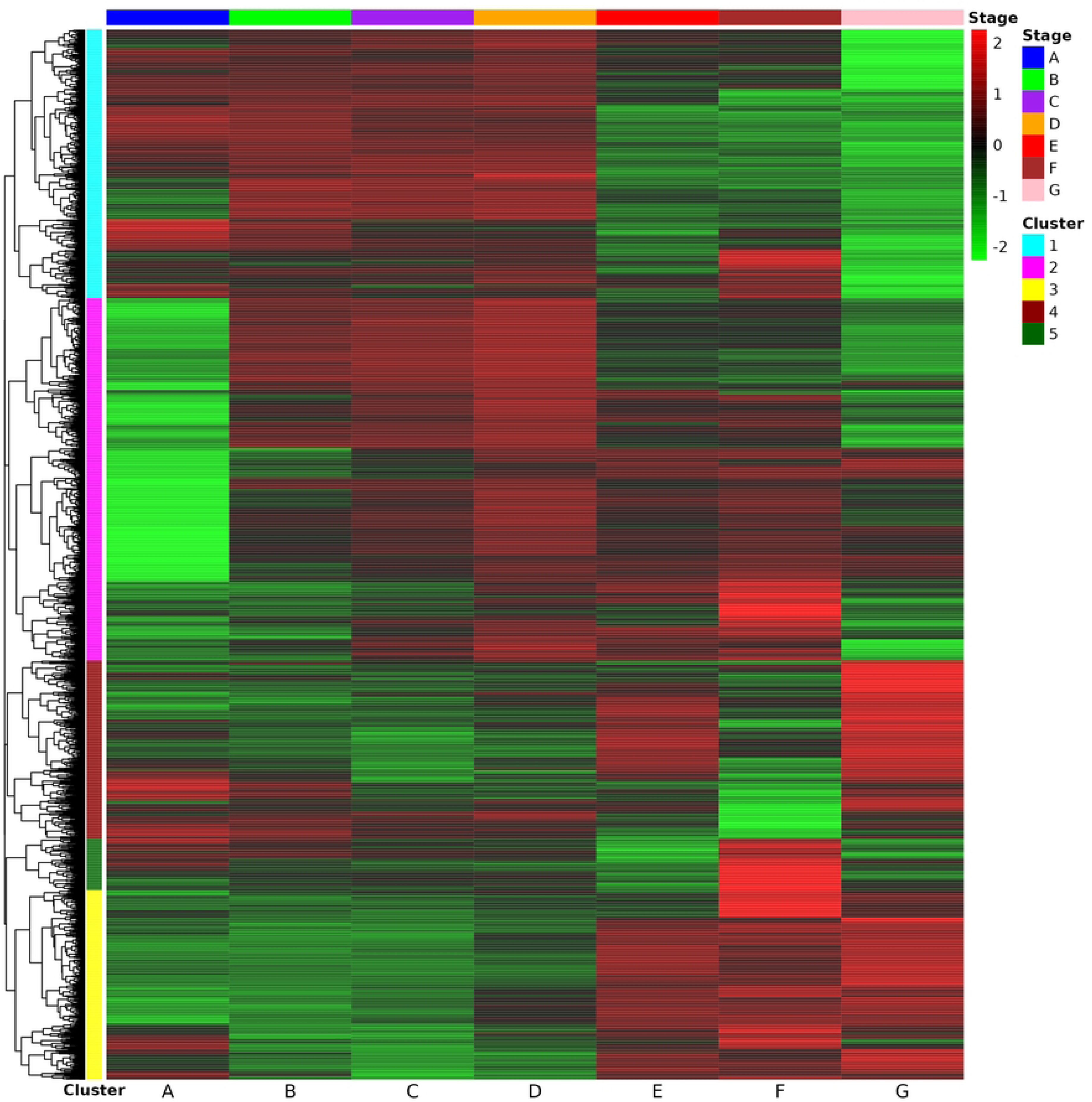
DEGs estimated by pairwise comparison of the seven stages.

Clustering analysis was performed for all 3234 DEGs based on expression and is shown in a heat map (Fig 4A). The analysis classified the DEGs into five groups (1 to 5) exhibiting different gene expression patterns, with counts of 827, 1116, 584, 548 and 159 genes for clusters 1, 2, 3, 4, and 5, respectively. The difference in the temporal expression patterns suggests stage-specific roles of these grouped DEGs throughout seed development. Groups 1, 2 and 3 represented the two largest clusters (827, 116, and 584 DEGs respectively) while exhibiting contrasting expression patterns. The DEGs in cluster 1 displayed the predominant and highest expression levels at stages A - D, while those in cluster 3 displayed the predominant and highest expression at later stages of development (15-21 DAF). The DEGs in cluster 2 were the highest in number and showed a continuous decrease in gene expression at stage A (3 DAF) and expression increased as seeds matured with the highest expression at stage D (12 DAF).

**Fig 4.**
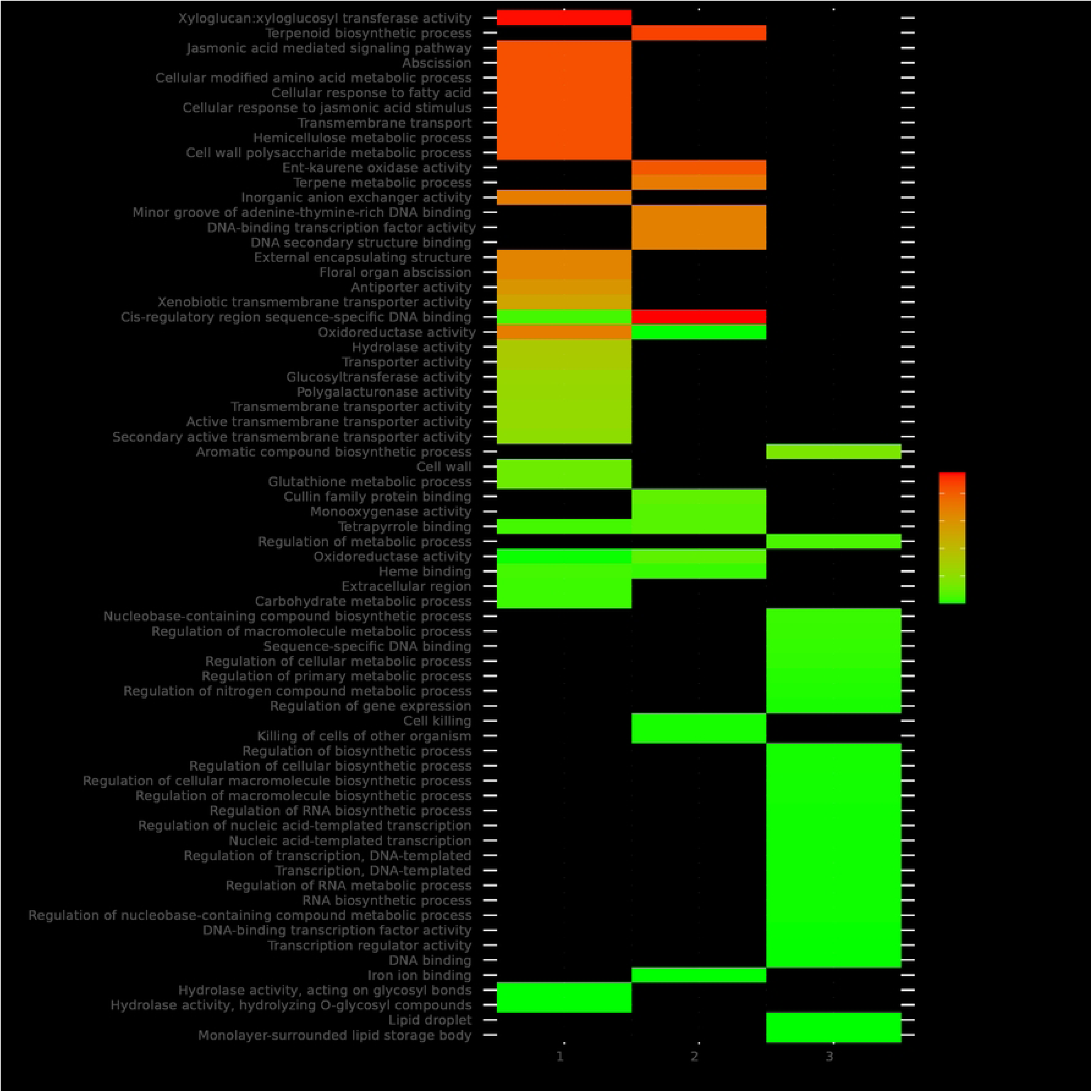
Clustering of differentially expressed genes across seven stages during seed development (A) Expression patterns of genes per cluster, The colours indicate the fold change with red = upregulated, black = regulated and green = downregulated genes (B) GO terms enriched for the larger clusters.

Enrichment analysis of cluster 1 showed that A-D-specific genes were enriched for the jasmonic acid-mediated signalling pathway (GO:0009867), Abscission (GO:0009838) and metabolic processes (GO:0010383, GO:0006575, GO:0005975) and the enriched biological processes occurred in the cell wall (Fig 4B). This stage represents the earlier stage of seed development. In comparison, cluster 3 showed that the stage E-G (15-21 DAF) genes were enriched for regulation of the RNA transcription process (GO:2001141), cellular metabolic process (GO:0019222) and DNA binding (GO:0003677). Despite the high number of DEGs in cluster 2, they showed fewer enriched categories, with the highest enriched category being Terpenoid biosynthesis (GO:0016114), Oxidoreductase activity (GO:0016491), and DNA-binding (GO:0000981)

GO annotation of these differentially expressed genes (DEGs) identified major biological processes, including metabolism, regulation of biological activities, cell wall organization, photosynthesis, and DNA replication (Table 3S). Top KEGG pathways enriched included carbon metabolism, plant hormone signal transduction, biosynthesis of cofactors, and glycogenesis (Table 4S).

### Analysis of key genes involved in seed development

We examined the differential expression of previously reported domestication-related genes [41] and genes associated with seed proteins [17] [5] [42] across the seven stages of lupin seed development, revealing significant variations (Fig 5 and Table 5S). Genes encoding storage proteins, such as cupin and Protease inhibitor family proteins, most displaying peak upregulation at stages E and G (15 and 21 DAF) (Fig 5a and b). Similarly, expression of three phytohormones related genes, AUX/IAA showed downregulated at the early stages of development, and upregulation at the later stage (Fig 5d) while a gene encoding gibberellins (GAs) was high at the later stage (21 DAF) of development (Fig 5b). Eleven genes encoding lipoxygenase showed regulation during seed development, with Lupan_010321 showing upregulation throughout the developmental stages. On the other hand, two protease inhibitors showed upregulated at stage Avs E and A vs G (15 and 21 DAF). Ten genes encoding YABBY TFs showed upregulation, mostly at later stages of seed development.

**Fig 5.**
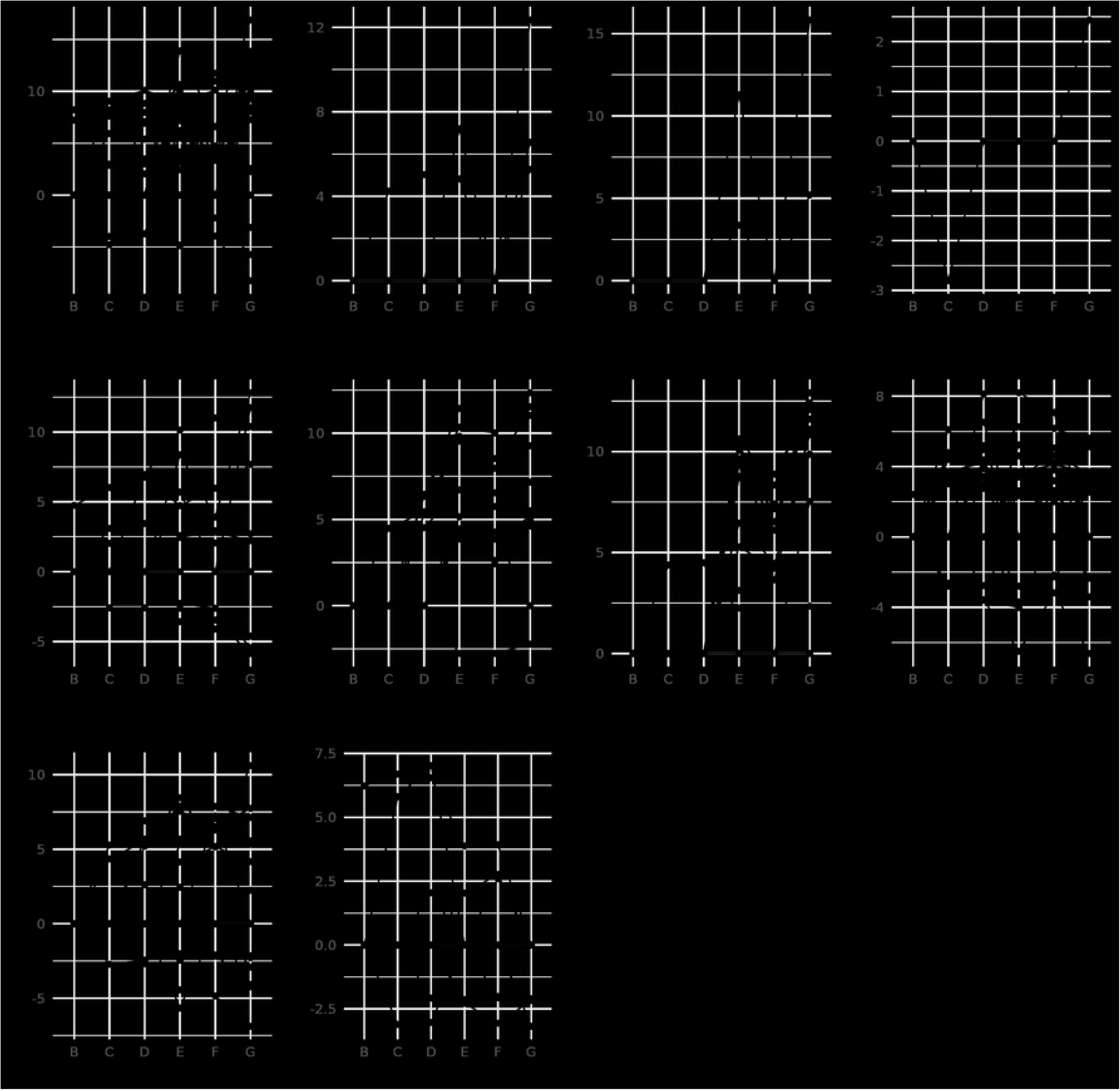
The transcription dynamics of genes involved in seed storage, phytohormone and domestication-related traits (a) Cupin; (b) Gibberellin; (c) Protease inhibitor; (d) AUX-IAA family; (e) Lipoxygenase; (f) YABBY; (g)Oleosin; (h) Alkaloid; (i) Vernalization; (j) Reduced shattering.

The analysis of expression patterns of domestication-related genes associated with reduced shattering showed diversity in expression. For instance, Lupan_005954 showed low regulation during early stages (3-12 DAF) and increased expression at 15-18 DAF, while Lupan_012033 exhibited higher expression at 18 DAF compared to other stages (Fig. 5j). Additionally, genes related to alkaloid metabolism exhibited diverse expression profiles. For instance, the gene associated with alkaloid synthesis, Lupan_032699, showed downregulation at stage C (12 DAF) and upregulation at stage G (21 DAF). In addition, Lupan_013564 showed the highest upregulation (LogFC>6) specifically at stages Cand D, representing 9-12 DAF (Fig. 5h).

### DEGs involved in TFs

Transcription factors (TFs) could participate in many aspects of cellular processes in seed development in seed development, thus the expression dynamics of TF genes involved in lupin seed development were investigated. According to the RNA-seq data, a total of 2486 genes belonging to 47 TF families were identified as differentially expressed in at least one developmental stage (Fig.7). The top 10 largest families were MYB (235), bHLH (235), C2H2(177), ERF (154), NAC (136), WRKY (117), LBD (110), HD-ZIP (98), Dof (84), and bZIP (84). The comparison among these three developmental stages found that 89 (79 up and 10 down-regulated), 282 (188 up and 94 down-regulated), 371 (274 up and 97 down-regulated), 511 (362 up and 149 down-regulated), 616 (440 up and 176 down-regulated), and 619 (390 up and 229 down-regulated) TFs had significant differential expression in A vs B, A vs C, A vs D, A vs E, A vs F, A vs G, respectively. Among these, the TFs with the largest number were M-Type-MADS (12), MYB (31), MYB (39), bHLH (56), bHLH (60), bHLH (67) for these seven stages respectively (Table S6). The BES1, GeBP, GRF, RAV, Nin-like, LFY and S1Fa-like families were upregulated in 18-21 DAF only suggesting their functions may be involved in the late stage of seed development or negatively during the grain maturation. Higher numbers of WRKY and NAC families showed upregulation at a mid-later stage of development (15-21 DAF) indicating their involvement in grain filling and maturation.

**Fig 6.**
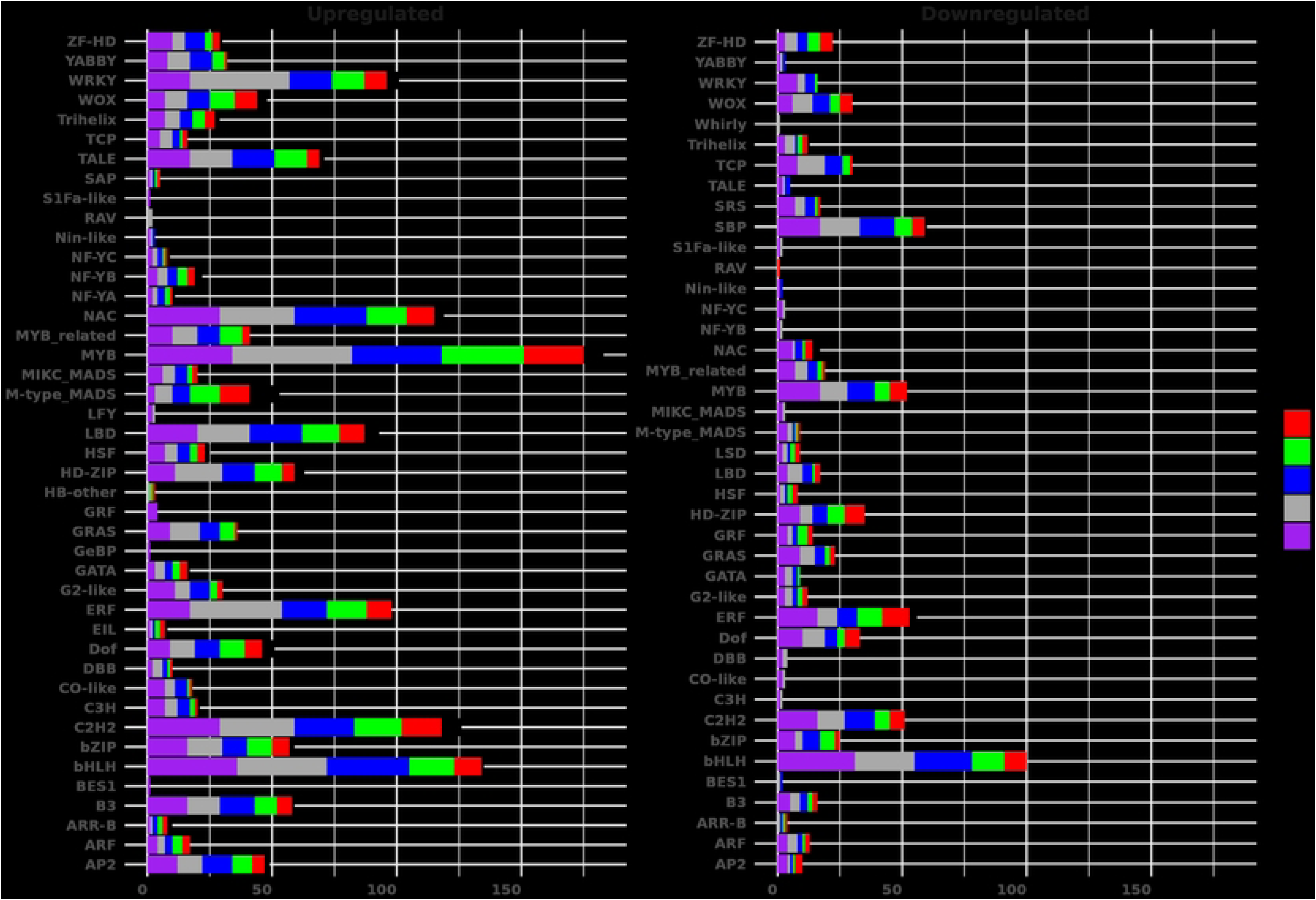
Counts of transcription factor family differentially expressed genes during seed development.

### Validation of DEGs by qRT-PCR

Based on the results mentioned above, 8 genes with significantly differential expression during seed development were selected for validation by performing qRT-PCR assays to validate the correlation against RNA-seq. The expression of these DEGs was consistent with the RNA-seq data results, confirming the transcriptome data’s reproducibility (Fig S2).

## Discussion

The development of seeds in crop plants significantly influences both seed fate and the ultimate yield and quality of the seeds. Therefore, a comprehensive understanding of the molecular mechanisms underlying seed development is essential for enhancing seed yield and quality in crop plants. Numerous genes involved in seed development have been identified and functionally validated, with molecular pathways elucidated in model plants such as Arabidopsis and Medicago [13, 31]. Additionally, transcriptome analyses in various crop plants have provided insights into the transcriptional dynamics of seed development and have suggested potential developmental pathways [6, 9, 43, 44, 44, 45]. In this study, we conducted a transcriptome analysis of lupin seed during development to gain insights into its transcriptional dynamics and identify candidate genes potentially involved in seed development.

The analysis of gene expression levels across different stages of lupin seed development (3, 6, 9, 12, 15, 18, and 21 days after flowering) revealed fluctuations in abundance and variance of these genes indicating their dynamic regulatory role throughout seed development. Cytochrome P450’s dramatic rise in later stages indicates its involvement in secondary metabolism and defence mechanisms during seed maturation [46]. The genes encoding Annexin repeat and lipoxygenase show high early-stage expression, declining towards the later stages, which suggests their roles in early seed development and initial cellular responses. Lipoxygenase has been reported to play crucial role in defence responses against abiotic stresses [47] while Annexins have been reported to play a role in the Golgi-mediated secretion of newly synthesized plasma membrane and wall material in plant cells [48]. These results indicate a temporal variation in transcript levels for many highly expressed genes during seed development, likely mirroring the underlying morphological and metabolic changes occurring throughout the seed development process.

GO term enrichment analysis of the highly abundant genes during lupin seeds development closely mirrors that of other dicotyledonous seeds, with an initial phase of cell division, followed by cell expansion, and culminating in maturation. The enrichment revealed significant enrichment in categories such as catalytic activity, binding, cellular processes, and metabolic processes, as well as cellular components including the cell and membrane. These results indicate that seed development is tightly regulated by genetic programming, which is likely associated with increased metabolic and catalytic activities within cellular and membrane structures. This was in agreement with earlier studies on chickpea [6], soybean [10], peanut [9], pigeon pea [49], *Medicago* [14] and Arabidopsis [13]. This enhanced understanding of lupin seed development provides valuable insights into the genetic and molecular mechanisms that underlie this critical process.

The expression of most common genes was predominantly observed during the early stages, indicating a progressive reduction in transcriptome dynamics as seeds advanced in development. This reduction may be attributed, at least partially, to the cessation of numerous transcriptional activities as the seeds advance toward maturity [50]. Early stages exhibited heightened metabolic activity, characterized by significant protein turnover, including both synthesis and degradation. In contrast, later stages were distinguished by activities related to protein storage and nutrient reservoir formation. Furthermore, the analysis revealed that most common genes were expressed throughout all stages of seed development, with only 1.61 % showing differential expression at various stages. This observation aligns with previous studies on soybean [12], chickpea [6], peanut [9], pigeon pea [49] and *Arabidopsis* [13], which suggested that genes involved in seed functions are largely consistent across developmental stages, despite potential variations in transcriptional activity. However, each developmental stage may possess a small, specific set of genes unique to that stage [51].

In the early stages of development, genes including that encoding glutathione transferase, alpha/beta hydrolase fold enzymes, thioredoxin-like proteins, purine nucleobase transporters, aluminium-activated malate transporters (ALMT), glycosyl hydrolase family 5 cellulases, leucine-rich repeat (LRR) proteins, the YABBY protein, glutathione transferase, and ML domain proteins were upregulated. Based on these findings, it is plausible to speculate on the possible roles of these gene families during lupin seed development. Alpha/beta hydrolase fold enzymes have been reported to be involved in lipid metabolism, providing energy and structural components crucial for seed growth in rice [52]. Thioredoxin-like proteins likely play a role in protecting developing seeds from oxidative stress in *Medicago* [53], thereby ensuring proper seed maturation, while purine nucleobase transporters may facilitate nucleic acid synthesis and energy metabolism, supporting rapid cell division during seed development [54]. LRR proteins may participate in developmental processes and defence responses, while ML domain proteins could be involved in lipid signalling and pathogen recognition, safeguarding seed development in lupin [55, 56].

We observed the differential expression patterns of seed storage protein genes encoding cupin, which are known to play crucial roles in plant development and defence [57]. These genes were called conglutins, are divided into four groups α, β, γ and δ, and are encoded by small gene families, respectively: *ALPHA1*-*3*, *BETA1-7*, *GAMMA1-2* and *DELTA1-4* [5, 58]. Previous studies on this group of cupin-encoded genes have shown variations in RNA expression during seed development in different lupin species [58, 59, 60]. In our current study, genes belonging to three of the main seed storage proteins, α, β and δ showed expression during lupin seed development, confirming their role as seed proteins [5]. In addition, the expression pattern of cupin genes encoding β and α conglutins, was previously reported to increase in transcription activities at 30 DAF [59]. These findings are supported by our current study, where four genes encoding β-conglutins (BETA1, BETA 2, BETA 4, BETA 6), three α conglutins (ALPHA1, ALPHA2, ALPHA3) and a DELTA2 showed upregulation at 15-21 DAF. These findings indicate the role of cupin genes in grain filling and could be targeted for manipulating seed protein for nutrition and optimal growth. Of note is the pattern of expression of seed storage protein, ALPHA2 (Lupan_012649), which showed downregulation at early stages and upregulation at later stages of seed development. Although ALPHA2 and ALPHA3 had been reported to be closely related to each other than to ALPHA1[58], whether this variation in expression is important in seed development remains to be determined. The similar expression pattern of these seed storage genes suggests their expression may be regulated by a common mechanism and there may be a master regulator (s) to ensure overall protein quantity within the seed is maintained. The expression of β-conglutins and α conglutins had been reported to increase at 15 DAF with high level of GA application but no increase in protein amounts [59], and in the current study, genes encoding gibberellin were found to be expressed in the same pattern as the seed storage protein. Consistent with this hypothesis are the results from silencing soybean β-conglycinin protein which caused an increase of glycinin [61], and the fine-tuning with post-transcriptional regulation of storage protein synthesis [62] and other environmental variations [63].

A high level of quinolizidine alkaloids (QAs) is a trait in many lupin species, as these chemical compounds protect plants from pests and fungi [64], however, alkaloids are major antinutritional factors and provide a bitter taste [65]. Previous studies showed QAs were synthesized in the aerial part of the plant, [66], but results are still divided on whether QAs are exclusively transported or partially synthesised *in situ* seed [67]. The gene encoding lysine/ornithine decarboxylase (LCD) is reported to catalyze the first step of quinolizidine alkaloid biosynthesis [68]. The expression of LCD has been confirmed to be associated with alkaloid content in *L. angustifolius* by several independent studies [69, 70]. In the current study, Lupan_020529 gene, which encodes LCD, was not differentially expressed and had a low mean transcript abundance of 69.4. Our study revealed the expression of genes linked to QA’s biosynthesis. Lupan_020878, encoding purine permease transporter 1 (PUP1) [71], which is generally involved in alkaloid biosynthesis and transport, and reported to have similar expression pattern to LCD gene [72], suggesting an increase in biosynthesis and transportation of alkaloid at the later stage of seed development. PUP1 had been reported previously as a potential quinolizidine alkaloid biosynthesis gene because it revealed a similar expression pattern as lysine/ornithine decarboxylase, which catalyzes the first step of quinolizidine alkaloid biosynthesis [72]. Other alkaloid-related genes that showed similar expression included genes encoding copper amine oxidase, NAD-dependent epimerase, and carboxylesterase-1. A gene expression study involving transcriptome sequencing of four accession and RT-PCR profiling of 14 accessions differing in alkaloids content found Lupan_13780 gene, encoding ETHYLENE *RESPONSIVE TRANSCRIPTION FACTOR RAP2-7* and a *cis*-regulated component Lupan_013784, encoding *Fe SUPEROXIDE DISMUTASE 2* to have opposite direction of the association. Although both genes were not differentially expressed in the current study, we found both genes’ mean transcript abundance of 3.96 and 2662.96 for Lupan_13780 and Lupan_013784, respectively.

Transcription factors (TFs) play essential roles as regulatory proteins in the modulation of gene expression, influencing nearly all vital processes in plant development [73] [13]. Given the dynamic nature of seed development, it is critical to examine the TFs involved. Our study revealed that most TFs associated with seed developmental processes exhibited high expression levels during the early stages, specifically in seed tissues at 3-12 days after flowering (DAF). Furthermore, many of the well-established TFs known for their roles in seed development (bHLH, AP2, MYB, ERF, C2H2, NAC, WRKY, and C3H zinc finger families) showed differential expression, thus reinforcing the validity of our findings. Almost all AP2 and HAP transcription factors (TFs) exhibited high expression levels during the early stages of seed development, whereas a subset of WRKY and C3H zinc finger TFs showed increased expression at later stages. This pattern supports the previous study that gene activity diminishes as seeds develop [74]. The preferential expression of these TFs in the early stages of lupin seed development implies their significant role in processes such as protein turnover, synthesis of storage compounds, and general metabolism, which are predominant during this period. Conversely, the limited expression of these TFs during later stages suggests that only a small fraction is involved in the induction of seed maturity, as gene activity substantially decreases as maturity approaches Similar transcriptional dynamics have been reported in other species, such as soybean and rice, highlighting conserved mechanisms of seed development across different plant taxa [30, 75]. The presence of TF families such as FAR1, LFY, NF-X1, HB-PHD, STAT, SAP, and HRT-like, which were expressed but not differentially regulated, suggests that these TFs may play more stable, constitutive roles during seed development. These TFs may play important roles in basal regulatory processes or could respond to stimuli outside the developmental windows examined in this study. Further investigation into this specific group of TFs beyond 21 DAF could provide valuable insights into the mechanisms underlying seed maturation and dormancy.

## Conclusion

In conclusion, we profiled the transcriptomes of developing seeds at seven representative stages and provide a global transcriptome view of the developmental processes of seed development. The identification of specific genes and their activities at different stages of seed development provides valuable information for scientists to understand how lupin seeds grow and develop, which can lead to better crop management practices. This transcriptome sequencing analysis indicated that the general pattern of seed development in lupin shares similarities with other dicot seeds. However, the preferential expression of specific genes or gene families may contribute to the unique nutritional properties of lupin seeds. Therefore, this study lays the groundwork for the characterization of candidate genes, offering novel insights into seed development and providing valuable resources for enhancing lupin breeding programs. Future research should focus on characterizing the specific roles of the candidate genes identified in this study to better understand their functions in lupin seed development and their potential applications in breeding programs.

## Abbreviations

CPM: Counts Per Million
DAF: Days after flowering
DEGs: Differentially expressed genes
FDR: False discovery rate
GO: Gene Ontology
LogFC: Log2 fold change
MF: Molecular function
TF: Transcription factor
TS: Tissue Specific

## Authors contributions

Conceptualization: L.G, V.W, and K.M.; sample collection: N.F, H.C, V.W and L.G.; RNA sample preparation for sequencing: V.W, H.C, and N.F; data curation and analysis: V.W, R.T, H.T, M.A, M.M, K.M. Conducting the qPCR validation experiments and analysis; writing original draft preparation: V.W. All authors reviewed and edited the final manuscript.

## Supporting information

**S1 Fig:** (A) Highlights genes with the highest average expression levels and (B) Genes with the greatest variability in expression.

**S2 Fig**: Validation of RNA-seq data using RT-qPCR

**S1 Table:** Highly abundant genes expressed during seed development at 3, 6, 9, 12, 15, 18, and 21 days after flowering (DAF), corresponding to stages A, B, C, D, E, F, and G of lupin seed development.

**S2 Table:** Gene Ontology (GO) enrichment analysis of highly abundant genes at 3, 6, 9, 12, 15, 18, and 21 days after flowering (DAF), corresponding to stages A, B, C, D, E, F, and G of lupin seed development.

**S3 Table:** GO enrichment analysis of DEGs during seed development at 3, 6, 9, 12, 15, 18, and 21 days after flowering (DAF), corresponding to stages A, B, C, D, E, F, and G of lupin seed development.

**S4 Table:** Kyoto Encyclopedia of Genes and Genomes (KEGG) pathway analysis of all DEGs during seed development at 3, 6, 9, 12, 15, 18, and 21 days after flowering (DAF), corresponding to stages A, B, C, D, E, F, and G of lupin seed development.

**S5 Table:** Differentially expressed genes and their annotations of seed specific genes during seed development at 3, 6, 9, 12, 15, 18, and 21 days after flowering (DAF), corresponding to stages A, B, C, D, E, F, and G of lupin seed development. Genes with P. Adj. < 0.05 and LogFC ≥ 2 are upregulated, while those with LogFC ≤ -2 are downregulated.

**S6 Table:** Transcriptional family and their genes distribution at different stages of development

---

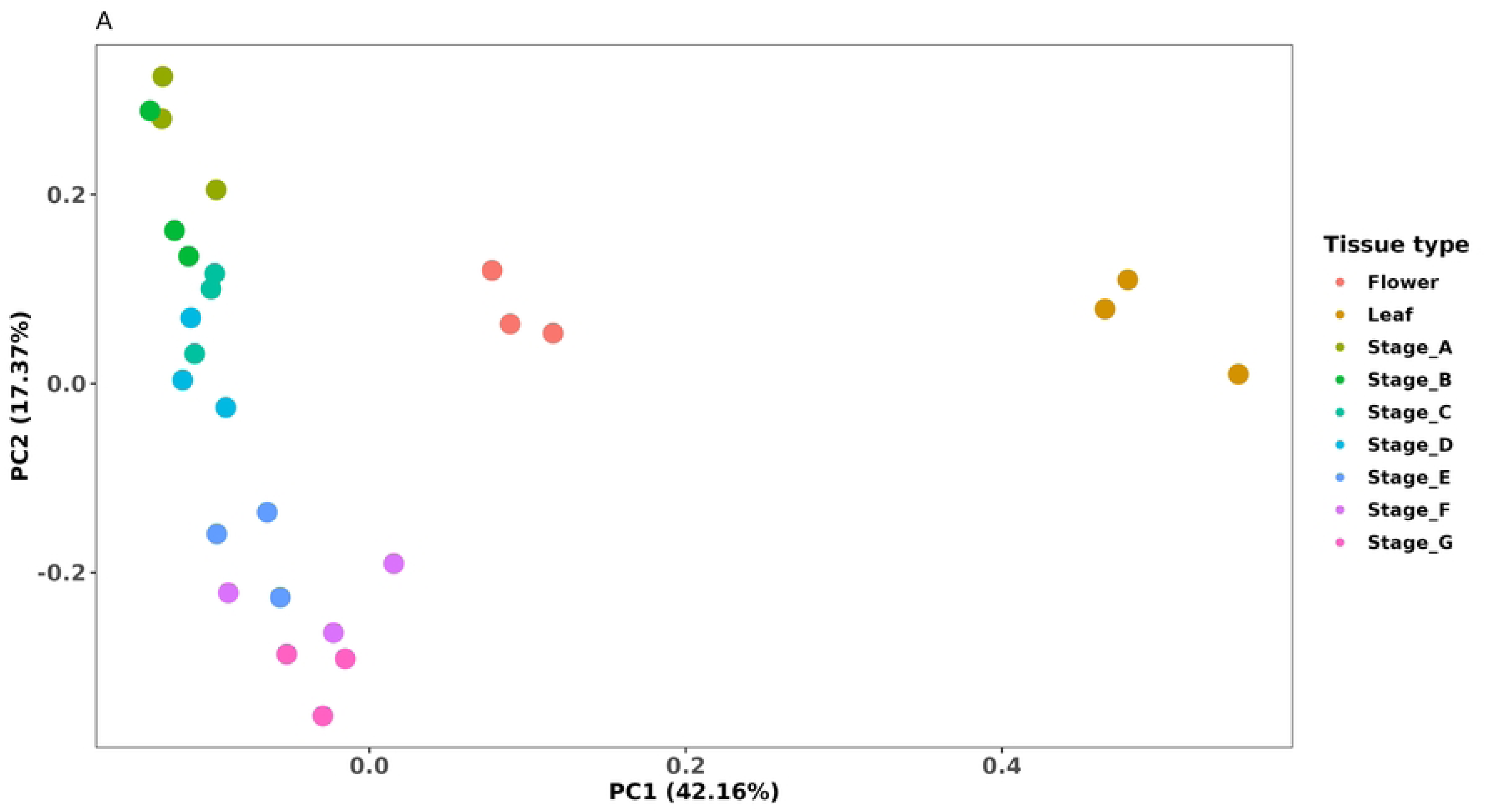

## References

1. Danan Gu, Kirill Andreev, Matthew E. Dupre. Major Trends in Population Growth Around the World[J]. China CDC Weekly. 2021; 604–613. doi:10.46234/ccdcw2021.160

2. Safdar LB, Foulkes MJ, Kleiner FH, Searle IR, Bhosale RA, Fisk ID, et al. Challenges facing sustainable protein production: Opportunities for cereals. Plant Communications. 2023;4: 100716. doi:10.1016/j.xplc.2023.100716

3. Vishnyakova M, Kushnareva A, Shelenga T, Egorova G. Alkaloids of narrow-leaved lupine as a factor determining alternative ways of the crop’s utilization and breeding. VAVILOVSKII ZHURNAL GENETIKI I SELEKTSII. 2020;24: 625–635. doi:10.18699/VJ20.656

4. Islam S, Yan G, Appels R, Ma W. Comparative proteome analysis of seed storage and allergenic proteins among four narrow-leafed lupin cultivars. Food Chem. 20120527th ed. 2012;135: 1230–8. doi:10.1016/j.foodchem.2012.05.081

5. Foley RC, Jimenez-Lopez JC, Kamphuis LG, Hane JK, Melser S, Singh KB. Analysis of conglutin seed storage proteins across lupin species using transcriptomic, protein and comparative genomic approaches. BMC Plant Biology. 2015;15: 106. doi:10.1186/s12870-015-0485-6

6. Pradhan S, Bandhiwal N, Shah N, Kant C, Gaur R, Bhatia S. Global transcriptome analysis of developing chickpea (Cicer arietinum L.) seeds. Frontiers in Plant Science. 2014;5. doi:10.3389/fpls.2014.00698

7. Yang L, Yang L, Ding Y, Chen Y, Liu N, Zhou X, et al. Global Transcriptome and Co-Expression Network Analyses Revealed Hub Genes Controlling Seed Size/Weight and/or Oil Content in Peanut. PLANTS-BASEL. 2023;12. doi:10.3390/plants12173144

8. Garg R, Singh VK, Rajkumar MS, Kumar V, Jain M. Global transcriptome and coexpression network analyses reveal cultivar-specific molecular signatures associated with seed development and seed size/weight determination in chickpea. Plant J. 2017;91: 1088–1107. doi:10.1111/tpj.13621

9. Guo F, Zhu X, Zhao C, Zhao S, Pan J, Zhao Y, et al. Transcriptome Analysis and Gene Expression Profiling of the Peanut Small Seed Mutant Identified Genes Involved in Seed Size Control. Int J Mol Sci. 20220827th ed. 2022;23. doi:10.3390/ijms23179726

10. Asakura T, Tamura T, Terauchi K, Narikawa T, Yagasaki K, Ishimaru Y, et al. Global gene expression profiles in developing soybean seeds. Plant Physiology and Biochemistry. 2012;52: 147–153. doi:10.1016/j.plaphy.2011.12.007

11. Yang S, Miao L, He J, Zhang K, Li Y, Junyi G. Dynamic Transcriptome Changes Related to Oil Accumulation in Developing Soybean Seeds. International Journal of Molecular Sciences. 2019;20: 2202. doi:10.3390/ijms20092202

12. Severin AJ, Woody JL, Bolon Y-T, Joseph B, Diers BW, Farmer AD, et al. RNA-Seq Atlas of Glycine max: A guide to the soybean transcriptome. BMC Plant Biology. 2010;10: 160. doi:10.1186/1471-2229-10-160

13. Le BH, Cheng C, Bui AQ, Wagmaister JA, Henry KF, Pelletier J, et al. Global analysis of gene activity during Arabidopsis seed development and identification of seed-specific transcription factors. Proceedings of the National Academy of Sciences. 2010;107: 8063–8070. doi:10.1073/pnas.1003530107

14. Gallardo K, Firnhaber C, Zuber H, Héricher D, Belghazi M, Henry C, et al. A Combined Proteome and Transcriptome Analysis of Developing Medicago truncatula Seeds: Evidence for Metabolic Specialization of Maternal and Filial Tissues*. Molecular & Cellular Proteomics. 2007;6: 2165– 2179. doi:10.1074/mcp.M700171-MCP200

15. Dam S, Laursen BS, Ørnfelt JH, Jochimsen B, Stærfeldt HH, Friis C, et al. The Proteome of Seed Development in the Model Legume Lotus japonicus. Plant Physiology. 2009;149: 1325–1340. doi:10.1104/pp.108.133405

16. Borisjuk L, Rolletschek H, Radchuk R, Weschke W, Wobus U, Weber H. Seed Development and Differentiation: A Role for Metabolic Regulation. Plant Biology. 2004;6: 375–386. doi:10.1055/s-2004-817908

17. Kamphuis LG, Hane JK, Nelson MN, Gao L, Atkins CA, Singh KB. Transcriptome sequencing of different narrow-leafed lupin tissue types provides a comprehensive uni-gene assembly and extensive gene-based molecular markers. Plant Biotechnol J. 20140724th ed. 2015;13: 14–25. doi:10.1111/pbi.12229

18. Garg G, Kamphuis LG, Bayer PE, Kaur P, Dudchenko O, Taylor CM, et al. A pan-genome and chromosome-length reference genome of narrow-leafed lupin (Lupinus angustifolius) reveals genomic diversity and insights into key industry and biological traits. Plant Journal. 2022;111: 1252–1266. doi:10.1111/tpj.15885

19. Yu B, Gao P, Song J, Yang H, Qin L, Yu X, et al. Spatiotemporal transcriptomics and metabolic profiling provide insights into gene regulatory networks during lentil seed development. The Plant Journal. 2023;115: 253–274. doi:10.1111/tpj.16205

20. Liu H, Liang X, Lu Q, Li H, Liu H, Li S, et al. Global transcriptome analysis of subterranean pod and seed in peanut (Arachis hypogaea L.) unravels the complexity of fruit development under dark condition. Scientific Reports. 2020;10: 13050. doi:10.1038/s41598-020-69943-7

21. Verma SK, Mittal S, Wankhede DP, Parida SK, Chattopadhyay D, Prasad G, et al. Transcriptome Analysis Reveals Key Pathways and Candidate Genes Controlling Seed Development and Size in Ricebean (Vigna umbellata). Frontiers in Genetics. 2022;12. doi:10.3389/fgene.2021.791355

22. Jiang LZ, Liu CY, Fan Y, Wu Q, Ye XL, Li Q, et al. Dynamic transcriptome analysis suggests the key genes regulating seed development and filling in Tartary buckwheat (Fagopyrum tataricum Garetn.). Frontiers in Genetics. 2022;13. doi:10.3389/fgene.2022.990412

23. Wang Y, Nie L, Ma J, Zhou B, Han X, Cheng J, et al. Transcriptomic Variations and Network Hubs Controlling Seed Size and Weight During Maize Seed Development. Frontiers in Plant Science. 2022;13. Available: https://www.frontiersin.org/articles/10.3389/fpls.2022.828923

24. Mathur S, Paritosh K, Tandon R, Pental D, Pradhan AK. Comparative Analysis of Seed Transcriptome and Coexpression Analysis Reveal Candidate Genes for Enhancing Seed Size/Weight in Brassica juncea. Frontiers in Genetics. 2022;13. doi:10.3389/fgene.2022.814486

25. Zhao L, Li M, Ma X, Luo D, Zhou Q, Liu W, et al. Transcriptome analysis and identification of abscisic acid and gibberellin-related genes during seed development of alfalfa (Medicago sativa L.). BMC Genomics. 2022;23: 651. doi:10.1186/s12864-022-08875-0

26. D’Agostino N, Li W, Wang D. High-throughput transcriptomics. Scientific Reports. 2022;12: 20313. doi:10.1038/s41598-022-23985-1

27. Wang Z, Gerstein M, Snyder M. RNA-Seq: a revolutionary tool for transcriptomics. Nature Reviews Genetics. 2009;10: 57–63. doi:10.1038/nrg2484

28. Xiang D, Quilichini TD, Liu Z, Gao P, Pan Y, Li Q, et al. The Transcriptional Landscape of Polyploid Wheats and Their Diploid Ancestors during Embryogenesis and Grain Development. The Plant Cell. 2019;31: 2888–2911. doi:10.1105/tpc.19.00397

29. Hasan S, Furtado A, Henry R. Gene Expression in the Developing Seed of Wild and Domesticated Rice. International Journal of Molecular Sciences. 2022;23. doi:10.3390/ijms232113351

30. Zhang H, Hu Z, Yang Y, Liu X, Lv H, Song B-H, et al. Transcriptome profiling reveals the spatial-temporal dynamics of gene expression essential for soybean seed development. BMC Genomics. 2021;22: 453. doi:10.1186/s12864-021-07783-z

31. Verdier J, Kakar K, Gallardo K, Le Signor C, Aubert G, Schlereth A, et al. Gene expression profiling of M. truncatula transcription factors identifies putative regulators of grain legume seed filling. Plant Molecular Biology. 2008;67: 567–580. doi:10.1007/s11103-008-9320-x

32. Chen S, Zhou Y, Chen Y, Gu J. fastp: an ultra-fast all-in-one FASTQ preprocessor. Bioinformatics. 2018;34: i884–i890. doi:10.1093/bioinformatics/bty560

33. Dobin A, Davis CA, Schlesinger F, Drenkow J, Zaleski C, Jha S, et al. STAR: ultrafast universal RNA-seq aligner. Bioinformatics. 2013;29: 15–21. doi:10.1093/bioinformatics/bts635

34. Love MI, Huber W, Anders S. Moderated estimation of fold change and dispersion for RNA-seq data with DESeq2. Genome Biol. 2014;15: 550. doi:10.1186/s13059-014-0550-8

35. R Core Team R. R: A language and environment for statistical computing. 2013.

36. Huang DW, Sherman BT, Lempicki RA. Systematic and integrative analysis of large gene lists using DAVID bioinformatics resources. Nature Protocols. 2009;4: 44–57. doi:10.1038/nprot.2008.211

37. Kanehisa M, Goto S. KEGG: Kyoto Encyclopedia of Genes and Genomes. Nucleic Acids Research. 2000;28: 27–30. doi:10.1093/nar/28.1.27

38. Kanehisa M, Sato Y, Kawashima M. KEGG mapping tools for uncovering hidden features in biological data. Protein Science. 2022;31: 47–53. doi:10.1002/pro.4172

39. Jin J, Zhang H, Kong L, Gao G, Luo J. PlantTFDB 3.0: a portal for the functional and evolutionary study of plant transcription factors. Nucleic Acids Research. 2014;42: D1182–D1187. doi:10.1093/nar/gkt1016

40. Livak KJ, Schmittgen TD. Analysis of relative gene expression data using real-time quantitative PCR and the 2(-Delta Delta C(T)) Method. Methods. 2001;25: 402–8. doi:10.1006/meth.2001.1262

41. Plewiński P, Książkiewicz M, Rychel-Bielska S, Rudy E, Wolko B. Candidate Domestication-Related Genes Revealed by Expression Quantitative Trait Loci Mapping of Narrow-Leafed Lupin (Lupinus angustifolius L.). Int J Mol Sci. 20191112th ed. 2019;20. doi:10.3390/ijms20225670

42. Kamphuis LG, Foley RC, Frick KM, Garg G, Singh KB. Transcriptome Resources Paving the Way for Lupin Crop Improvement. In: Singh KB, Kamphuis LG, Nelson MN, editors. Lupin Genome. 2020. pp. 53–71. doi:10.1007/978-3-030-21270-4_5

43. Wang Y, Nie L, Ma J, Zhou B, Han X, Cheng J, et al. Transcriptomic Variations and Network Hubs Controlling Seed Size and Weight During Maize Seed Development. FRONTIERS IN PLANT SCIENCE. 2022;13. doi:10.3389/fpls.2022.828923

44. Yu Y, Zhu D, Ma C, Cao H, Wang Y, Xu Y, et al. Transcriptome analysis reveals key differentially expressed genes involved in wheat grain development. The Crop Journal. 2016;4: 92–106. doi:10.1016/j.cj.2016.01.006

45. Geng X, Dong N, Wang Y, Li G, Wang L, Guo X, et al. RNA-seq transcriptome analysis of the immature seeds of two Brassica napus lines with extremely different thousand-seed weight to identify the candidate genes related to seed weight. PLoS One. 20180130th ed. 2018;13: e0191297. doi:10.1371/journal.pone.0191297

46. Schuler MA, Werck-Reichhart D. Functional Genomics of P450S. Annual Review of Plant Biology. Annual Reviews; 2003. pp. 629–667. 10.1146/annurev.arplant.54.031902.134840

47. Rosahl S. Lipoxygenases in Plants -Their Role in Development and Stress Response. Zeitschrift für Naturforschung C. 1996;51: 123–138. doi:doi:10.1515/znc-1996-3-401

48. Wei H, Chen J, Zhang X, Lu Z, Lian B, Liu G, et al. Comprehensive analysis of annexin gene family and its expression in response to branching architecture and salt stress in crape myrtle. BMC Plant Biology. 2024;24: 78. doi:10.1186/s12870-024-04748-8

49. Parida SK, Pazhamala LT, Agarwal G, Bajaj P, Kumar V, Kulshreshtha A, et al. Deciphering transcriptional programming during pod and seed development using RNA-Seq in Pigeonpea (Cajanus cajan). PLoS ONE. 2016;11. doi:10.1371/journal.pone.0164959

50. Considine MJ, Considine JA. On the language and physiology of dormancy and quiescence in plants. Journal of Experimental Botany. 2016;67: 3189–3203. doi:10.1093/jxb/erw138

51. Sreenivasulu N, Wobus U. Seed-Development Programs: A Systems Biology–Based Comparison Between Dicots and Monocots. Annual Review of Plant Biology. Annual Reviews; 2013. pp. 189–217. 10.1146/annurev-arplant-050312-120215

52. Dolui AK, Vijayaraj P. Functional Omics Identifies Serine Hydrolases That Mobilize Storage Lipids during Rice Seed Germination. Plant Physiology. 2020;184: 693–708. doi:10.1104/pp.20.00268

53. Alkhalfioui F, Renard M, Vensel WH, Wong J, Tanaka CK, Hurkman WJ, et al. Thioredoxin-Linked Proteins Are Reduced during Germination of Medicago truncatula Seeds. Plant Physiology. 2007;144: 1559–1579. doi:10.1104/pp.107.098103

54. Andriotis VME, Smith AM. The plastidial pentose phosphate pathway is essential for postglobular embryo development in Arabidopsis. Proceedings of the National Academy of Sciences. 2019;116: 15297–15306. doi:10.1073/pnas.1908556116

55. Figueroa-Macías JP, García YC, Núñez M, Díaz K, Olea AF, Espinoza L. Plant Growth-Defense Trade-Offs: Molecular Processes Leading to Physiological Changes. International Journal of Molecular Sciences. 2021;22. doi:10.3390/ijms22020693

56. Shao Q, Liu X, Su T, Ma C, Wang P. New Insights Into the Role of Seed Oil Body Proteins in Metabolism and Plant Development. Frontiers in Plant Science. 2019;10. Available: https://www.frontiersin.org/journals/plant-science/articles/10.3389/fpls.2019.01568

57. Dunwell JM, Culham A, Carter CE, Sosa-Aguirre CR, Goodenough PW. Evolution of functional diversity in the cupin superfamily. Trends in Biochemical Sciences. 2001;26: 740–746. doi:10.1016/S0968-0004(01)01981-8

58. Foley RC, Gao LL, Spriggs A, Soo LY, Goggin DE, Smith PM, et al. Identification and characterisation of seed storage protein transcripts from Lupinus angustifolius. BMC Plant Biol. 20110404th ed. 2011;11: 59. doi:10.1186/1471-2229-11-59

59. Klajn N, Kapczyńska K, Pasikowski P, Glazińska P, Kugiel H, Kęsy J, et al. Regulatory Effects of ABA and GA on the Expression of Conglutin Genes and LAFL Network Genes in Yellow Lupine (Lupinus luteus L.) Seeds. International Journal of Molecular Sciences. 2023;24. doi:10.3390/ijms241512380

60. Duranti M, Consonni A, Magni C, Sessa F, Scarafoni A. The major proteins of lupin seed: Characterisation and molecular properties for use as functional and nutraceutical ingredients. Trends in Food Science & Technology. 2008;19: 624–633. doi:10.1016/j.tifs.2008.07.002

61. Kinney AJ, Jung R, Herman EM. Cosuppression of the α Subunits of β-Conglycinin in Transgenic Soybean Seeds Induces the Formation of Endoplasmic Reticulum–Derived Protein Bodies. The Plant Cell. 2001;13: 1165–1178. doi:10.1105/tpc.13.5.1165

62. Tabe L, Hagan N, Higgins TJV. Plasticity of seed protein composition in response to nitrogen and sulfur availability. Current Opinion in Plant Biology. 2002;5: 212–217. doi:10.1016/S1369-5266(02)00252-2

63. Bourgeois M, Jacquin F, Savois V, Sommerer N, Labas V, Henry C, et al. Dissecting the proteome of pea mature seeds reveals the phenotypic plasticity of seed protein composition. PROTEOMICS. 2009;9: 254–271. doi:10.1002/pmic.200700903

64. Wink M, Meißner C, Witte L. Patterns of quinolizidine alkaloids in 56 species of the genus Lupinus. Phytochemistry. 1995;38: 139–153. doi:10.1016/0031-9422(95)91890-D

65. Bassoli A, Borgonovo G, Busnelli G. Alkaloids and the Bitter Taste. Modern Alkaloids. 2007. pp. 53–72. doi:10.1002/9783527621071.ch3

66. Bunsupa S, Yamazaki M, Saito K. Quinolizidine alkaloid biosynthesis: recent advances and future prospects. Frontiers in Plant Science. 2012;3. Available: https://www.frontiersin.org/journals/plant-science/articles/10.3389/fpls.2012.00239

67. Lee MJ, Pate JS, Harris DJ, Atkins CA. Synthesis, transport and accumulation of quinolizidine alkaloids in Lupinus albus L. and L. angustifolius L. Journal of Experimental Botany. 2007;58: 935– 946. doi:10.1093/jxb/erl254

68. Bunsupa S, Katayama K, Ikeura E, Oikawa A, Toyooka K, Saito K, et al. Lysine Decarboxylase Catalyzes the First Step of Quinolizidine Alkaloid Biosynthesis and Coevolved with Alkaloid Production in Leguminosae. The Plant Cell. 2012;24: 1202–1216. doi:10.1105/tpc.112.095885

69. Frick KM, Foley RC, Kamphuis LG, Siddique KHM, Garg G, Singh KB. Characterization of the genetic factors affecting quinolizidine alkaloid biosynthesis and its response to abiotic stress in narrow-leafed lupin (Lupinus angustifolius L.). Plant Cell Environ. 20180510th ed. 2018;41: 2155–2168. doi:10.1111/pce.13172

70. Kroc M, Koczyk G, Kamel KA, Czepiel K, Fedorowicz-Strońska O, Krajewski P, et al. Transcriptome-derived investigation of biosynthesis of quinolizidine alkaloids in narrow-leafed lupin (Lupinus angustifolius L.) highlights candidate genes linked to iucundus locus. Sci Rep. 20190219th ed. 2019;9: 2231. doi:10.1038/s41598-018-37701-5

71. Berger JD, Buirchell BJ, Luckett DJ, Nelson MN. Domestication bottlenecks limit genetic diversity and constrain adaptation in narrow-leafed lupin (Lupinus angustifolius L.). Theor Appl Genet. 20111103rd ed. 2012;124: 637–52. doi:10.1007/s00122-011-1736-z

72. Yang T, Nagy I, Mancinotti D, Otterbach SL, Andersen TB, Motawia MS, et al. Transcript profiling of a bitter variety of narrow-leafed lupin to discover alkaloid biosynthetic genes. J Exp Bot. 2017;68: 5527–5537. doi:10.1093/jxb/erx362

73. Guo Z, Maki M, Ding R, Yang Y, zhang B, Xiong L. Genome-wide survey of tissue-specific microRNA and transcription factor regulatory networks in 12 tissues. Scientific Reports. 2014;4: 5150. doi:10.1038/srep05150

74. Sreenivasulu N, Wobus U. Seed-development programs: a systems biology-based comparison between dicots and monocots. Annual review of plant biology. 2013;64: 189–217.

75. Fait A, Angelovici R, Less H, Ohad I, Urbanczyk-Wochniak E, Fernie AR, et al. Arabidopsis seed development and germination is associated with temporally distinct metabolic switches. Plant Physiol. 2006/09/08 ed. 2006;142: 839–854. doi:10.1104/pp.106.086694

